# Autophagy and evasion of immune system by SARS-CoV-2. Structural features of the Non-structural protein 6 from Wild Type and Omicron viral strains interacting with a model lipid bilayer. ^†^

**DOI:** 10.1101/2022.01.05.475107

**Authors:** Emmanuelle Bignon, Marco Marazzi, Stéphanie Grandemange, Antonio Monari

**Affiliations:** Université de Lorraine and CNRS, LPCT UMR 7019, F-54000 Nancy France; Department of Analytical Chemistry, Physical Chemistry and Chemical Engineering and Chemical Research Institute “Andres M. del Rio” (IQAR), Universidad de Alcalá, 28805 Alcalá de Hénares, Spain; Université de Lorraine and CNRS, CRAN UMR 7039, F-54000 Nancy France; Université de Paris and CNRS, ITODYS, F-75006 Paris, France

## Abstract

The viral cycle of SARS-CoV-2 is based on a complex interplay with the cellular machinery, which is mediated by specific proteins eluding or hijacking the cellular defense mechanisms. Among the complex pathways called by the viral infection autophagy is particularly crucial and is strongly influenced by the action of the non-structural protein 6 (Nsp6) interacting with the endoplasmic reticulum membrane. Importantly, differently from other non-structural proteins Nsp6 is mutated in the recently emerged Omicron variant, suggesting a possible different role of autophagy. In this contribution we explore, for the first time, the structural property of Nsp6 thanks to long-time scale molecular dynamic simulations and machine learning analysis, identifying the interaction patterns with the lipid membrane. We also show how the mutation brought by the Omicron variant may indeed modify some of the specific interactions, and more particularly help anchoring the viral protein to the lipid bilayer interface.

**Electronic Supplementary Information (ESI) available:** Analysis protein of the secondary structure and of the specific lipid/amino acid interactions. RMSF per amino acid. Distribution of the distance between the center of mass of the 89 to 99 α-helix and the center of the lipid bilayer. Analysis of the behavior of the 195 to 207 α-helix. See DOI: 10.1039/x0xx00000x

## Introduction

The emergence in late 2019 of a novel positive sense, single-stranded RNA β-coronavirus, shortly afterwards named SARS-CoV-2, has led to the outbreak of a pandemic, COVID-19, which has since then severely affected every country worldwide. Indeed, and despite its relatively low mortality ratio, the high transmissibility of SARS-CoV-2,^1–3^ coupled to the potential development of serious outcomes requiring intensive care treatment,^4^ has resulted in a considerable strain posed on the health systems and has led to serious social distancing and containment measures, including full lockdowns. If the development and the large-scale deployment, at least in Western countries, of efficient vaccines,^5–7^ including the revolutionary mRNA strategy,^8–11^ have allowed a much better containment of the pandemic and a significant decrease in both deaths and intensive care admissions, SARS-CoV-2 remains a considerable and not fully mastered threat. Indeed, the year 2021 has been characterized by the emergence of different SARS-CoV-2 variant, classified as variant of concern (VOC) by the World Health Organization (WHO), which due to their higher transmissibility have rapidly become dominant, replacing the original viral strains, even if the mutation rate of SARS-CoV-2 is much slower compared to other RNA viruses. Also thank to the presence of an exonuclease acting as a proof reading during viral genome replication, its wide diffusion is, for obvious statistical reasons, prone to favorize dominant mutations. After the emergence first of Alpha^12^ and Beta^13^ (beginning of 2021) and Delta variant^14^ (summer 2021) a novel strain, styled Omicron and accumulating a high density of point mutations has been reported in Southern African countries the 25^th^ of November 2021.^15^ The Omicron variant,^16–19^ also called B.1.1.529, is characterized by both a higher transmissibility and the partial capacity to infect subjects with prior immunity obtained either via vaccination or precedent infection.^20^ Despite some preliminary data seem to point to a less severity rate of Omicron compared to the original variant,^21^ its high transmissibility and the partial evasion of precedent immunization constitutes a drastic problem.^22^

From a molecular biology point of view,^23,24^ SARS-CoV-2 genome is constituted by a large, ∼ 30 k bases, positive-sense single-stranded RNA fragment, which is enveloped in a membrane virion. After cellular infection the viral genome is translated into two large polyproteins, PP1 and PP2, and some structural proteins, such as spike (S, responsible for the interaction with the cellular receptor and the membrane fusion, and concentrating most of the Omicron mutations),^25^ nuclear (N) and envelope (E) proteins. In turn the original PPs are self-cleaved by two proteases giving rise to the so-called non-structural proteins (Nsp), which are responsible for crucial viral processes related to its replication and resistance to the host\ immune system. Indeed, among the non-structural proteins one should cite, in addition to the proteases, the RNA-dependent RNA polymerase^28,29^ which is responsible for the genome replication, via a temporary negative-stranded RNA template, the exonuclease complex,^30^ and the SARS-Unique Domain (SUD) which may sequester RNA to impede triggering apoptotic signals.^31,32^ Furthermore, membrane Nsps are also present and tend to accumulate in the endoplasmic reticulum (ER), i.e. the replication compartment of SARS-CoV-2. Nsp6 has also been recognized as capable of interfering with Type I interferon pathway by blocking the activation of Tank binding kinase allowing viral evasion of innate immune response.^33^ Nsp6 has also been reported as an important regulator of autophagy in infected cells.^34,35^ As a matter of fact, coronaviruses’ infected cells present a higher number of autophagosomes, the latter being much smaller than in non-infected cells, a phenomenon in which the Nsp6 protein plays a crucial role.^34^ These different roles of NSP6 reflect the complex equilibrium between immune response and viral replication.^36^ Interfering with different steps of autophagy favors the viral survival and propagation.^37^ Indeed, while autophagosomes incorporating exogenous and endogenous protein material may actively participate in the elimination/destruction of viral components and hence enhance adaptative immunity by delivering viral antigens, they may also form protective compartment in which viral replication can take place using autophagic products such as metabolites and substrates. Reducing the size of the autophagosome may, thus, hamper the fusion of autophagosomes with lysosomes to prevent the elimination of the viral material while maintaining the favorable environment for the viral replication and maturation.^35^

The structure of many key SARS-CoV-2 structural and non-structural proteins has been resolved,^28,38,39^ from the first day of the pandemic, and the relation between their structure and activity has since been also complemented by multiscale molecular modeling and simulation,^24,31,40–42^ also tackling enzymatic reactivity.^26,27^ Undoubtedly, the S protein, also in complex with the human ACE2 receptor, and the viral proteases have been the main target for structural biology and molecular modeling simulations.^43–47^ Of note, the proposition of some possible viral inhibitors has also been undertaken with some success. In contrast, the structure, and hence the mechanisms, of other Nsps including Nsp6,^33^ has been much less studied despite its fundamental biological role. Notably, while no experimentally resolved structure of membrane-embedded Nsp6 is not available, the combination of sequence homology and machine learning approach^48^ has allowed proposing a putative starting structure. In this contribution we aim at filling this gap validating the proposed Nsp6 structure through extended all atom molecular dynamic (MD) simulation, identifying the key structural motifs allowing an efficient interaction with a lipid bilayer. Our results are also coherent with the ones of Kumar et al. for the WT strain.^49^

The SARS-CoV-2 variants exhibit a high density of mutations, mainly concentrated on the spike coding sequence, which influence greatly the binding to the human ACE2 receptor and its viral invasion capacity.^50^ Interestingly, the Omicron variants also presents the deletion of three amino acids, namely L105, S106 and G107, from the Nsp6 sequence.^25^ The three deleted amino acids are located at the polar head/water interface where they connect, via a distorted loop, two trans-membrane α-helices. Their absence can clearly influence the protein/membrane interaction and hence have a non-negligible role in autophagy.^51^ Hence, in the following we provide MD simulations on the mutated protein to be compared with the one originated from the native strain.

Our results in addition to rationalizing the dynamical properties of Nsp6 and its interaction with lipid bilayers, our results also tackle the effects of the mutations of the Omicron variant in a crucial protein responsible for the virus maturation and immune system elusion.

## Results

As shown in Figure 1 the structure of the Wild Type (WT) Nsp6 presents all the key features of typical transmembrane proteins: transmembrane helices bundle and extramembranous connectors. Furthermore, the structure of the protein retrieved by machine learning approaches is found to be remarkably stable. Indeed, all along the 2 µs-long MD simulation of Nsp6 embedded in a lipid bilayer only very limited and local deviation from the initial structure are observed. This is evidenced by the time series of the root mean square deviation (RMSD) of the protein which after an equilibration period of about 200-330 ns exhibits an extended plateau to reach an average value of 2.88 ± 0.01 Å. Remarkably, even during the first equilibration period the maximum value of the RMSD barely exceeds 4 Å confirming the global stability of the protein structure and its favorable interaction with the lipid bilayer. Besides, the membrane structural parameters also dos not exhibit remarkable fluctuations during the production run, confirming the stability of the overall system – see Figure S1. Hence, our results can also be regarded as a first independent confirmation of the proposed structure of Nsp6. From the analysis of the protein secondary structure along the MD simulation we may identify some crucial features, which deserve discussion. As shown in Figure 1B and 1C we may identify 8 rather long transmembrane α-helices (TM1 to TM8), which are mainly composed by hydrophobic residues and which assume a clear transmembrane bundle structure, which is a widespread pattern in membrane proteins, see for instance rhodopsin and other receptors. Despite their slightly different length and some minor changes in their orientation with respect to the membrane axis, all the eight α-helices are remarkably stable both concerning their secondary and tertiary structure and exhibits only negligible displacements all along the simulation. Once again, the rigidity of the transmembrane core may be regarded as a typical feature of stable membrane proteins. On top of the transmembrane core one can also identify six shorter and more flexible α-helices and 2 β -sheets extending at the interface between the lipid polar head and water. Interestingly, these interfacial structures tend to position in parallel conformation with respect to the membrane plan, and the β-helices are disposed almost symmetrically on the two membrane/water interfaces.

**Figure 1.**
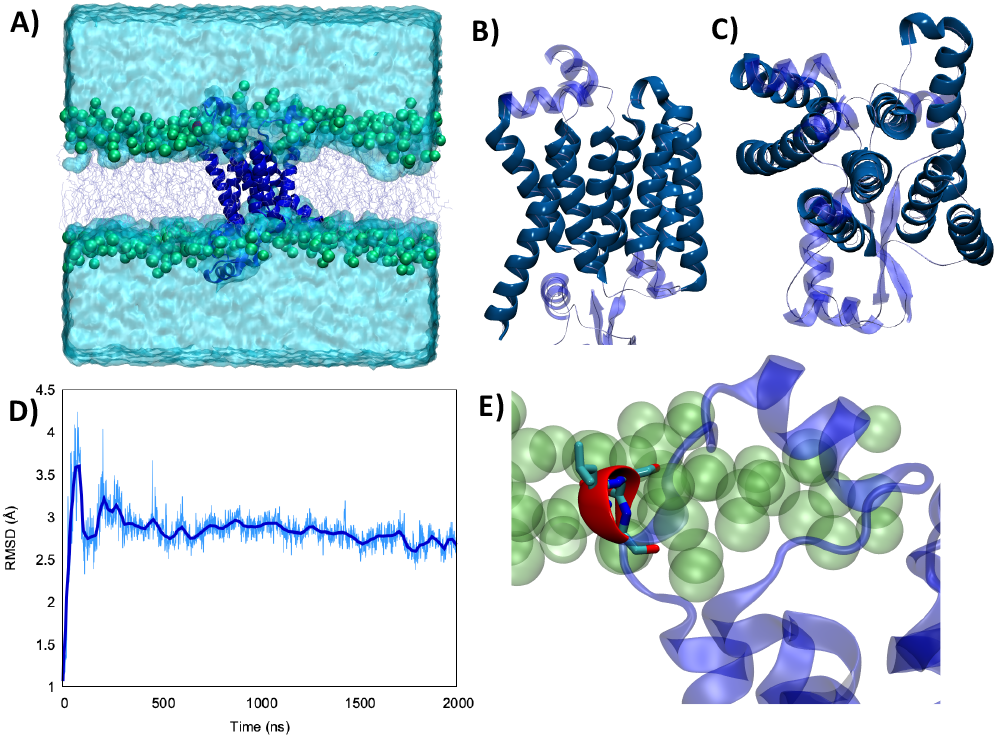
A) Representation of the simulation box showing the WT Nsp6 protein embedded in a model lipid bilayer surrounded by water buffer. Side (B) and top (C) view of the Nsp6 protein highlighting the secondary structure motifs. The transmembrane regions are represented in darker blue and opaque, while extramembranous areas are rendered in lighter blue and in transparence. D) Time series of the RMSD for Nsp6 along the MD simulation. E) Zoom on the three amino acids which are deleted in Omicron variant represented in red and licorice.

Differently from the transmembrane core structure these shorter motifs present a higher density of polar residue, coherently with their positioning at the polar head region. Furthermore, some transient electrostatic interactions, as well as hydrogen bonds, are formed between the phosphate or the choline moieties of the lipids and the interfacial amino acids. However, the interaction network is highly dynamic and evolves continuously all along the simulation without showing a dominant pattern. This fact may also partially contribute to justifying the higher flexibility of the extramembranous regions as compared to the core. These considerations are supported by an analysis of the secondary structure along the trajectory (Figure S2), showing the stability of the eight transmembrane helices and of the two isolated β-sheets, while all extramembranous α-helices are more flexible (especially between TM3 and TM4 and after TM8 toward the -C terminus), due to bending, turning and, more in general, the presence of non-structured short linkers. Of particular interest, see Figure 1E, is a very short α-helix composed of a triad of residues, namely L105, S106 and G107. Indeed, while the secondary structure is stable all along the MD simulation, this triad can be found inside an unstructured loop connecting the transmembrane core to the extramembranous α-helix formed by residues 89 to 99. This short helix is one of the most flexible and mobile moieties of Nsp6 and experiences significant oscillation on the membrane plan. Furthermore, L105, S106 and G107 are also the amino acids that are deleted from Nsp6 in the Omicron variant. Hence, this structural motif, and the nearby areas may experience the greater variability among the different strains.

For this reason, we performed an independent MD simulation of the Omicron Nsp6 variant, which has been obtained by manually mutating the WT strain. The results of the MD simulations are collected in Figure 2. From the analysis of the MD simulation it is apparent that the Omicron variant is still able to favorably interact with the lipid bilayer producing a stable aggregate (Figure 2A). Noteworthy, the structural parameters of the membrane (i.e., thickness, position of the membrane center on the Z axis and area per lipid) remain stable along the simulation for both variants (Figure S1). Furthermore, as evidenced by the time evolution of the RMSD, we may see that the global deformation of the protein is still rather small, and more importantly after a first equilibration region where it reaches 3.5/4.0 Å it stabilizes to an extended plateau at 3.0 Å. Coherently with what was observed for the WT Nsp6 while the transmembrane core is extremely rigid and experiences barely any fluctuation over the course of the MD simulation, the peripheral helices present a greater mobility and flexibility. Hence, the peripheral motifs will also show the higher difference between the WT and Omicron variant, as pictorially shown in Figure 2 C,D where representative snapshots for the two variants are superimposed. As expected, the peripheral α-helix comprised between the 89 and 99 residues, i.e. the region between TM3 and TM4, is indeed the one showing the larger deviation between the two structures (see Figure S2). As a matter of fact, the mutation in the Omicron strain involves the short loop connecting this helix to the protein core, hence justifying the higher variability of its structure. These global tendencies are also confirmed by the analysis of the flexibility profile at the residue level (Figure 2E) and by the root mean square fluctuation (RMSF, reported in Figure S3). Indeed, while most of the protein, and particularly the transmembrane core presents a very low flexibility, with the partial exception of TM7, which experiences considerable turning events in its -C terminal region (see Figure S2). As a matter of fact, two peaks can be observed corresponding to the residues 86-108 and 195-207 for the WT. These two regions represent two transmembrane helices protruding in the polar head region. Furthermore, the first flexible peak also encompasses the amino acid triad which is deleted in the Omicron variant. As for the comparison with the Omicron variant we may observe that while the rigidity profile is similar to the WT a rather larger flexibility is observed in correspondence of the two peaks. This may be related to the necessity of rearranging the local structure of the protein to account for the effects of the mutation.

**Figure 2.**
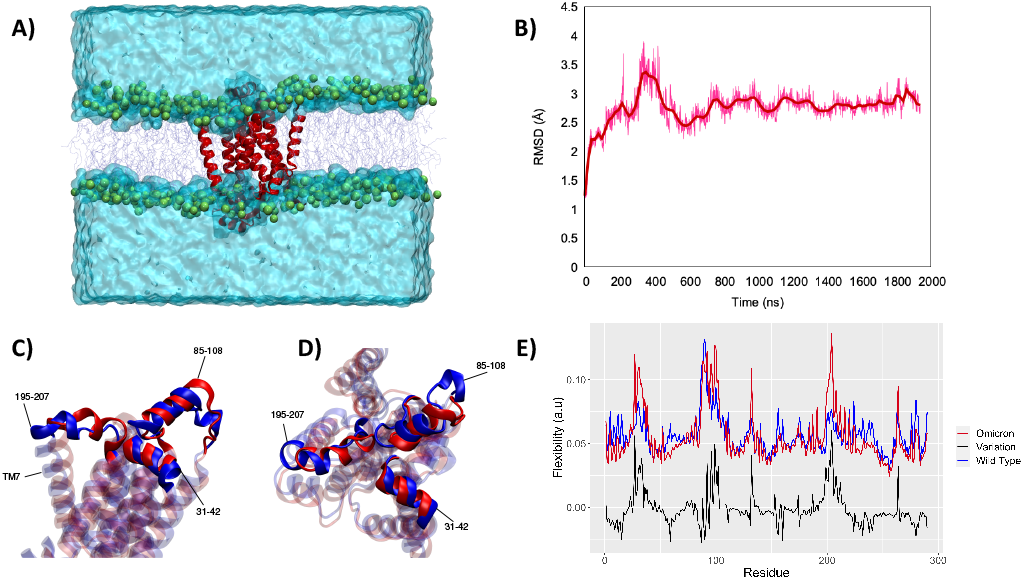
A) A snapshot of the full simulation box showing the stability of the interaction between Omicron Nsp6 and the lipid membrane. The structural stability of the latter is also revealed by the time series of the RMSD for the protein (B). Side (C) and top (D) view of the superposition of WT (blue) and Omicron (red) Nsp6, the extramembranous helices and the mobile transmembrane helix 7 (TM7) are evidenced. E) Per amino acid flexibility profile of the WT (blue) and Omicron (red) variants the point-to-point difference is also reported in black

To better analyze the global effects of the different flexibility and organization around the membrane we report in Figure 3 the evolution of the relative position of the center of mass of 89-99 helix with respect to the center of mass of the lipid bilayer for the two variants, evidencing a slightly different behavior of the two proteins. Indeed, concerning the projection of the distance along the (x,y) plan, i.e. the one parallel to the membrane, we may see that the WT peaks around the (−8.0,0.0) Å position while the Omicron variant concentrates at (−9.0,0.0) Å. More importantly, the 2D-distribution shows a different topology between the two variants; the Omicron strain evidences a bimodal distribution with the presence in addition to the main peak of a metastable state at (−13.0,-6.0) Å spanning for a couple of hundreds of ns. On the contrary the WT shows a more pronounced variability with the distribution of distances covering an extended region prior to its stabilization to the final stable state.

**Figure 3.**
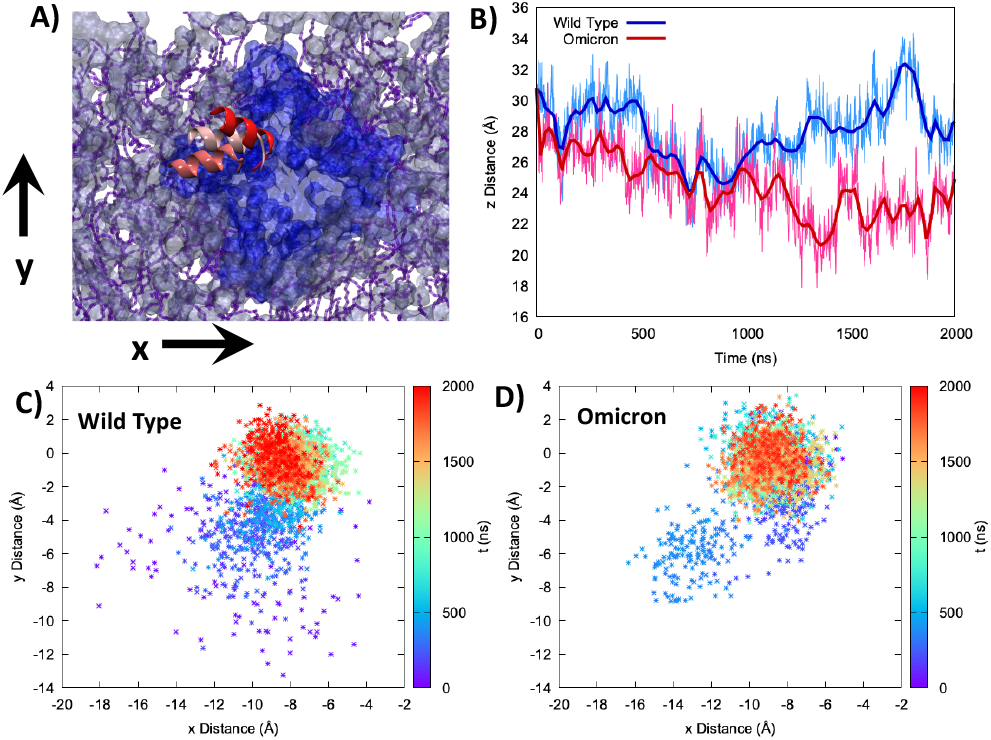
A) Representation of the mobility of the 89-99 α-helix by three different snapshots, shown with different reddish shades, are reported on top of the membrane and protein on the xy plane. B) Time series of the distance between the center of mass of the 85-108 helix and the lipid membrane projected on the z-axis parallel to the bilayer thickness for WT (blue) and Omicron (red). Projection on the xy plane of the distance between the center of mass of the 85-108 helix and the lipid membrane for the WT (C) and Omicron (D). The time series is given as a color map. The histograms of the distances distribution at various time interval are also reported in Figure S7.

The analysis of the projection of the distance on the z axis, i.e. the one perpendicular to the membrane, is perhaps even more significant and interesting. As shown by its time series reported in Figure 3B, we may evidence two very distinct behaviors between the two proteins. Indeed, in the case of the Omicron strain the peripheral helix is buried deeper inside the polar head region and assumes and average distance from the center of the bilayer of 24.8 ± 2.0 Å, to be compared with 27.7 ± 2.0 Å for the WT. Furthermore, even if less evident from the standard deviation, in the case of the WT we may evidence the oscillation between a buried (i.e. shorter distance) and an exposed (i.e. longer distance) conformation, which is translated in a rather broad distribution (see Figure S4). The oscillation period between the two WT conformations appears of the order of 400 ns, even though longer trajectories and a more extended sampling would be necessary to provide a quantitative estimation.

An analysis of the interactions between the extramembranous α-helix, containing the triad of residues deleted in the Omicron variant, and the surrounding membrane was performed by measuring the distance between the centers of mass of each residue and the center of mass of the polar heads found in a radius of 60 Å from the Leu105-Gly107 triad (Figure S5). As it can be seen, such deletion induced a consistent strengthening of the protein-membrane interaction, with an average distance that decreases from ca. 30 to ca. 15 Å, with charged Lys109 interacting at even shorter distances (peak at around 10 Å). Thus, we can state on a firm ground that the Omicron variant clearly favors a more buried conformation of this α-helix.

A similar behavior can also be observed for the 195-207 helix (Figure S6), which has also been identified as a flexibility hot-spot, and which is globally more buried in the case of the Omicron variant, presenting an average distance with the center of the membrane of 15.6 ± 1 Å compared to the 17.3 ± 2 Å observed for the WT. Furthermore, the projection of the distance on the (x,y) plan is also significantly different between the two variants with the WT peaking at (10.0,-1.0) Å and Omicron at (6.0,-1.5) Å as shown in Figure S6.

## Conclusions and Discussion

Nsp6 is a crucial protein in the SARS-Cov-2 viral cycle. In particular its capacity to modulate the autophagy response and the autophagosome topology is fundamental in regulating the delicate equilibrium between the establishment of favorable replication conditions and an efficient immune system response. However, its structure and its interaction with lipid bilayers has not been reported yet. In this contribution we fill this gap by validating the proposed Nsp6 structure using large-scale MD simulations and confirming the stability of the protein while identifying the crucial interaction pattern established with a model lipid bilayer.

Furthermore, the mutation of Nsp6 in the Omicron variant, consisting in the deletion of a triad of amino acids at the polar head interface, has been pinpointed as a potential concerning mutation due to its possibility of altering the modulation of autophagy by favoring the interaction with the membrane. Indeed, the molecular basis underlying autophagy will be due to the recruitment of lipid, specifically PIP2, by Nsp6 and the subsequent formation of the autophagosome vesicle. Hence, any mutation which would favor the interaction with the membrane is susceptible to alter this mechanism. While we have shown that the global structure of Nsp6 is not altered by the triad deletion, in particular concerning the transmembrane core, we have evidenced important modification of the structure and dynamic of the peripheral helices, as illustrated by the flexibility signature perturbation. More specifically, we have shown that the peripheral helices in the Omicron variant are more buried in the polar head region compared to the WT, hence suggesting stronger interaction and an increased capacity of recruiting specific lipids. Importantly, the structure of the POPC membrane does not show strong deviations between the two variants. Yet, further study about the complex effect of the membrane composition on its mechanical properties would provide a clearer picture of the interplay between the membrane structure and the biological aspects of the viral cycle. This would also include the study of the effects of lipids such as Phosphatidylinositol-4,5-bisphosphate (PIP2) which are known to play an important role in the autophagosome formation.^52^ However, while our results should be confirmed experimentally, they point to a different modulation of the autophagy between the two viral variants which may in part explain both the immune system resistance of the Omicron variant and its different pathological evolution. In the near future we plan to pursue this study on the one side including explicitly PIP2 in our simulations and on the other side experimentally comparing the induction of autophagy in cell lines in which WT or Omicron Nsp6 would have been expressed.

## Methodology

The initial proposed structure of SARS-CoV-2 Nsp6 has been retrieved from the Jumper et al. Deepmind web server^48^ collecting putative structures of COVID-19 related proteins obtained via the AlphaFold machine learning methodology^53^ fed by the sequences taken from the UniProt pre-release download.^54^ The structure of the Omicron variant has been manually constructed from the WT by deleting the L105, S106, G107 triad. The protocol used to construct the Omicron variant has been justified confirming by the AlphaFold-based^55^ structural prediction which confirms that the deletion of the aminoacid triad does not alter the global structure of the transmembrane α-helix bundle. Both WT and Omicron Nsp6 have been embedded in a lipid membrane bilayer constituted of two leaflets of 100 1-Palmitoyl-2-oleoylphosphatidylcholine (POPC) lipids, a water buffer has been added complemented with a physiological concentration of K+ and Cl-ions. Although cellular membranes are clearly more complex and present variable concentration of lipid species, including steroids, which can alter their rigidity, single lipid membranes may be considered as valid models of a biologically relevant lipid bilayers, and have already been used in different computational studies.^56,57^ The solvated membrane/protein structure has been prepared using the Charmm-gui web-server interface.^58^ The lipid and the protein have been represented by the Amber ff14^59,60^ force field while water has been modeled via TIP3P.^61^ MD simulations have been performed using the NAMD code,^62,63^ using periodic boundary condition (PBC) and the constant number of particle, pressure, and temperature (NPT) ensemble, using a Langevin thermostat and barostat.^64,65^ A time step of 4 fs has been used to integrate the Newton equation of motion thanks to the use of Hydrogen Mass Repartition (HMR)^66^ in combination with RATTLE and SHAKE.^67^ After minimization, thermalization and equilibration have been performed progressively releasing harmonic constraint on the protein backbone and the lipid during 40 ns, after which a 2 µs production run has been obtained for both the WT and the Omicron system. All the trajectories have been visualized and analyzed with the VMD code.^68^ Per residue Nsp6 flexibility profile was obtained by an in house machine learning protocol, based on the one proposed by Fleetwood and coworkers,^69^ considering the principal component analysis (PCA) of the protein based on the decomposition of the RMSD covariance matrix. In particular, we use the internal coordinates (inverse distance between geometric centers of two residues) along the trajectories as the input of PCA instead of the cartesian coordinates to provide better performance. The per residue importance (i.e. flexibility) was calculated by taking the sum of the weights of the principal components up to 80%, where the weight is defined as the eigenvalue of the corresponding principal component over the sum of all the eigenvalues of the RMSD covariance matrix. The secondary structure analysis of both WT and Omicron variant was performed by applying the Define Secondary Structure of Proteins (DSSP) algorithm,^70^ as implemented in the AmberTools21 suite of programs.^71^

## Author Contributions

Conceptualization, planning and data collection A.M. Data analysis and discussion A.M. E.B. M.M. S.G. All the authors participated to the draft realization and writing.

## Acknowledgements

Financial support from the French Higher Education research and Innovation Ministry (MESRI) and CNRS through the GAVO project is gratefully acknowledged. Calculations have been performed on LPCT and Itodys local computing clusters, on the regional Explor computing center under the project dancing under the light, and on the national GENCI center within the Seek&Destroy and Gavo allocations. AM thanks ANR (Agence Nationale de la Recherche) and CGI (Commissariat à l’Investissement d’Avenir) for their financial support of this work through Labex SEAM (Science and Engineering for Advanced Materials and devices) ANR 11 LABX 086, ANR 11 IDEX 05 02. The support of the IdEx “Université Paris 2019” ANR-18-IDEX-0001 and of the Platform P3MB is gratefully acknowledged. M.M. thanks the REACT-DISCoVER-UAH-CM (FEDER 2014 2020) funded by the Universidad de Alcalá, Comunidad de Madrid, and European Union, for financial support. Dr. Tao Jiang from Laboratoire de Chimie, ENS Lyon, is gratefully acknowledged for assistance with the PCA analysis and for providing the original script.

