## Supplementary Information for "Autophagy and evasion of immune system by SARS-CoV-2. Structural features of the Non-structural protein 6 from Wild Type and Omicron viral strains interacting with a model lipid bilayer. ^†^"

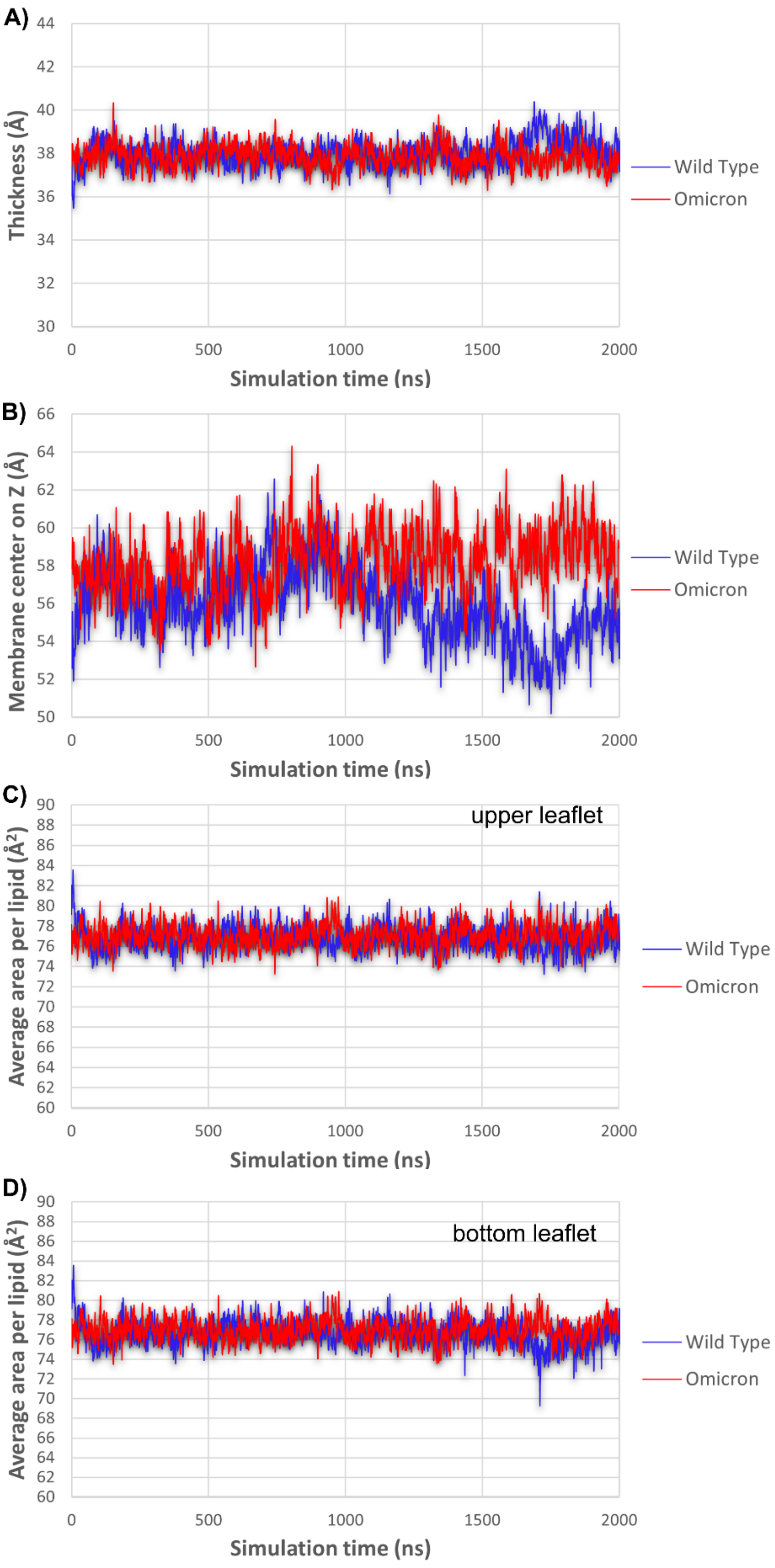


- **Figure S1.** Times series of the membrane parameters for the Wild Type NSP6 (red) and the Omicron variant (blue): A) Thickness, B) Position of the membrane center on the Z axis, and C) average area per lipid on the upper leaflet and D) the lower leaflet.

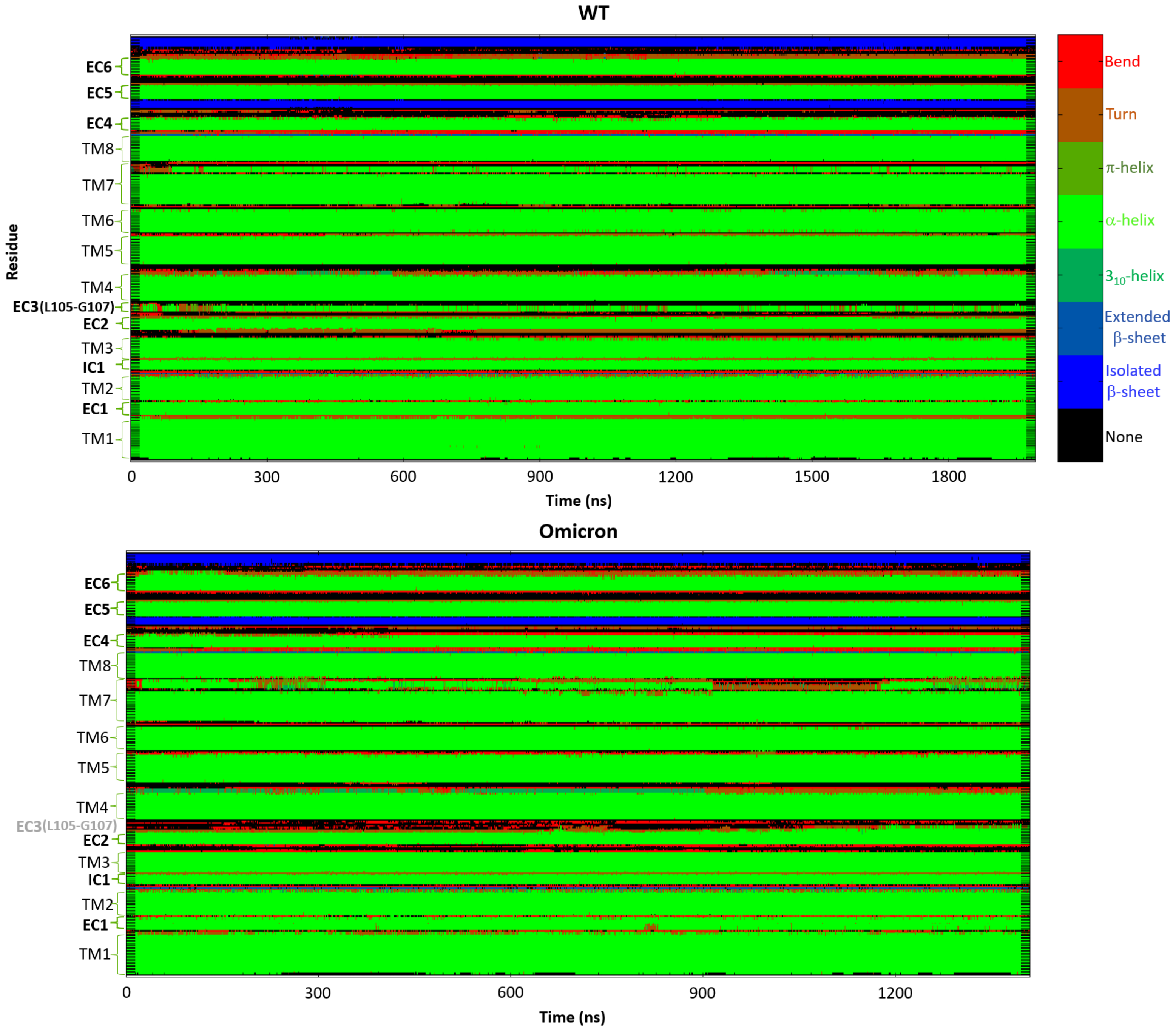


**Figure S2.** Secondary structure analysis of the WT (up) and Omicron (bottom) variant of the NSP6 protein, along each MD trajectory. The color code for each secondary structure is shown in the upper-right panel. The y-axis represents the primary structure from the -N to the -C terminus, including the depiction of all transmembrane α-helices (TM), extracellular side α-helices (EC), and intracellular side α-helices (IC). The residues' triad subjected to deletion is part of EC3.

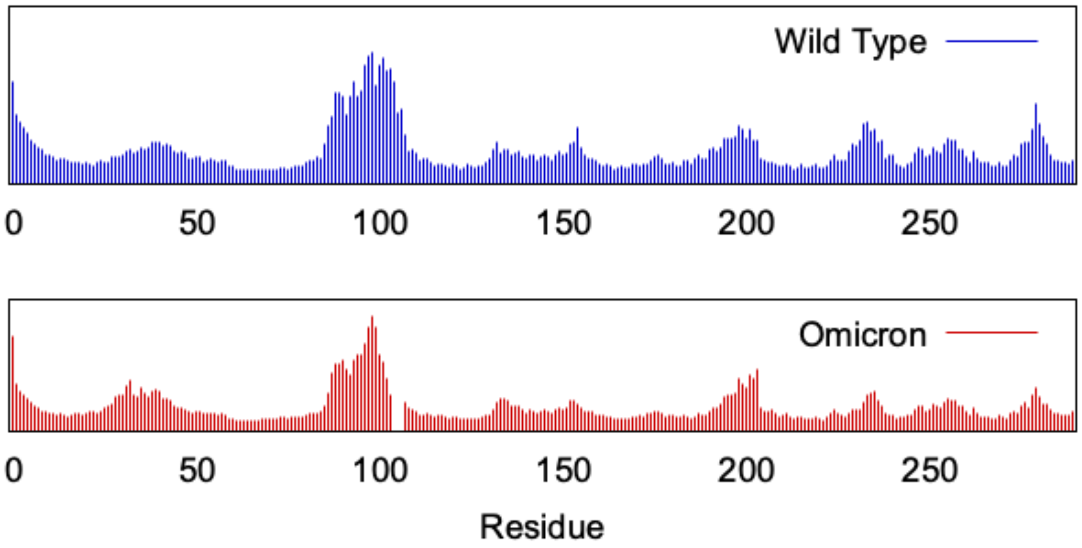


**Figure S3.** RMSF for the WT (top panel, blue) and Omicron (bottom panel, red) variant.

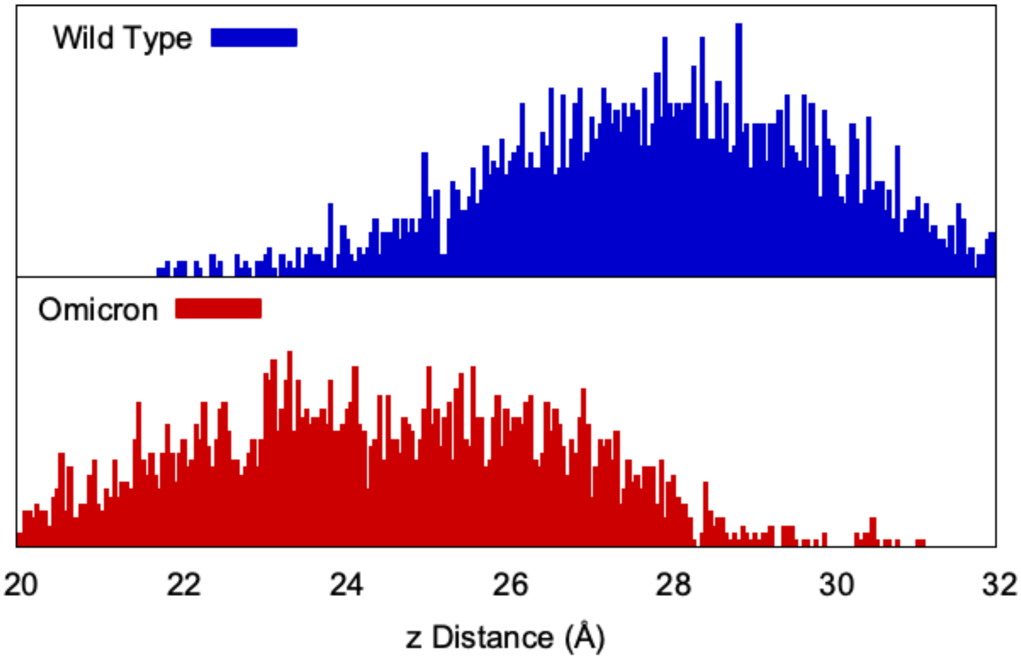


**Figure S4.** Distribution of the distance between the center of mass of the transmembranous 89-99 $\alpha$-helix and of the lipid bilayer for the WT (top panel blue) and Omicron variant (bottom panel, red).

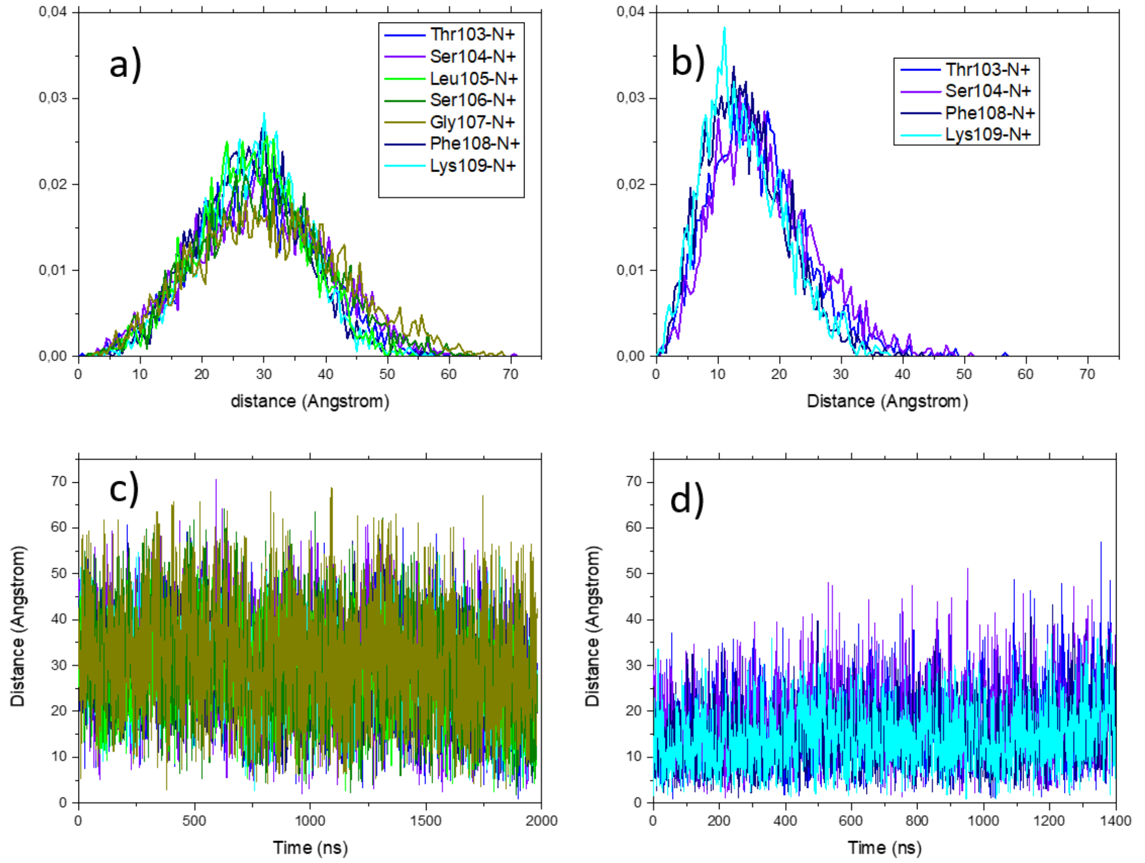


**Figure S5.** Distribution (a,b) and respective timeseries (c,d) of the distance between the center of mass of each considered residue and the center of mass of the polar heads of the 14 POPC residues found inside a radius of 60 Å form 105-107 triad, deleted in the Omicron variant.

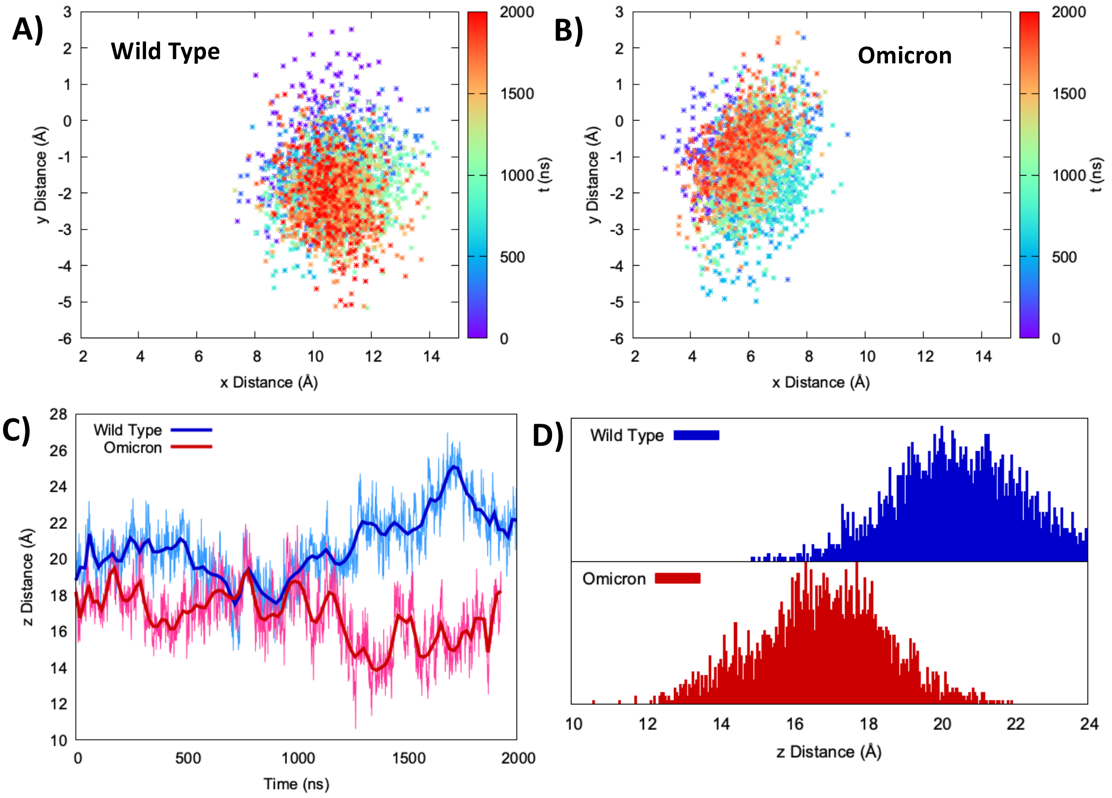


**Figure S6.** Distribution of the distances projected on the (x,y) between the center of mass of the 195 to 207 extramembranous $\alpha$-helix and the center of mass of the lipid bilayer for the WT (A) and Omicron variant (B). Time evolution (C) and distribution (D) of the distance between the center of mass of the 195 to 207 extramembranous $\alpha$-helix and the center of mass of the lipid bilayer projected on the z axis.

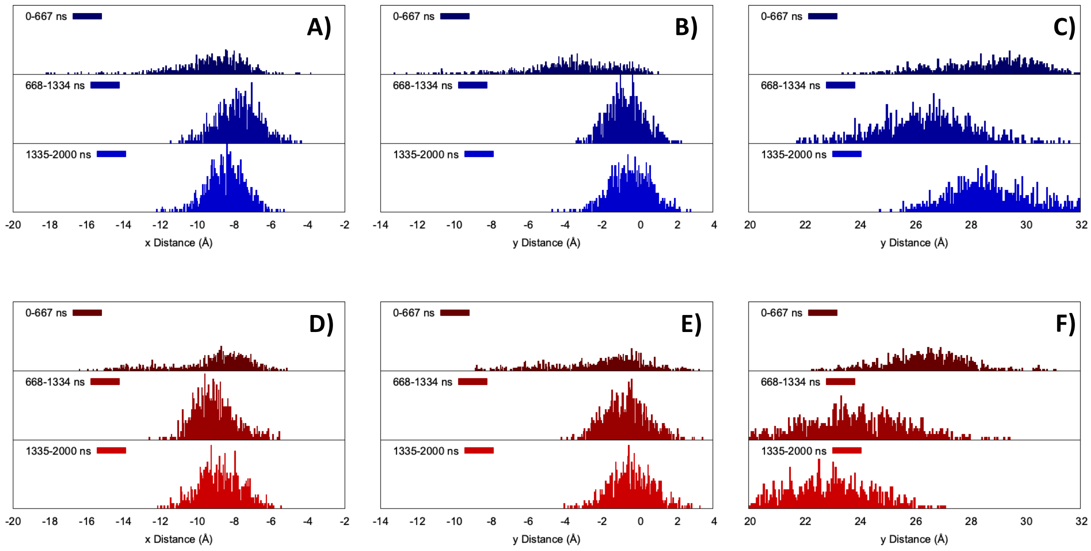


**Figure S7.** Distribution, at various time of the dynamics, of the distances projected on the x, y, and z axis (A, B, C, respectively) of between the center of mass of the 89 to 99 $\alpha$-helix and the center of mass of the lipid bilayer for the WT (A, B, C) and Omicron variant (D, E, F).
